# Moth resonant mechanics are tuned to wingbeat frequency and energetic demands

**DOI:** 10.1101/2024.01.30.578003

**Authors:** Ethan S. Wold, Brett Aiello, Manon Harris, Usama Bin Sikandar, James Lynch, Nick Gravish, Simon Sponberg

## Abstract

An insect’s wingbeat frequency is a critical determinant of its flight performance and varies by multiple orders of magnitude across Insecta. Despite potential energetic and kinematic benefits for an insect that matches its wingbeat frequency to its resonant frequency, recent work has shown that moths may operate off of their resonant peak. We hypothesized that across species, wingbeat frequency scales with resonance frequency to maintain favorable energetics, but with an offset in species that use frequency modulation as a means of flight control. The moth superfamily Bombycoidea is ideal for testing this hypothesis because their wingbeat frequencies vary across species by an order of magnitude, despite similar morphology and actuation. We used materials testing, high-speed videography, and a “spring-wing” model of resonant aerodynamics to determine how components of an insect’s flight apparatus (thoracic properties, wing inertia, muscle strain, and aerodynamics) vary with wingbeat frequency. We find that the resonant frequency of a moth correlates with wingbeat frequency, but resonance curve shape (described by the Weis-Fogh number) and peak location vary within the clade in a way that corresponds to frequency-dependent biomechanical demands. Our results demonstrate that a suite of adaptations in muscle, exoskeleton and wing drive variation in resonant mechanics, reflecting potential constraints on matching wingbeat and resonant frequencies.

## 1 Introduction

Fast cyclic movements are key to many organisms’ locomotor performance and have given rise to the convergent evolution of elastic structures that help offset inertial costs of locomotion by storing and releasing energy from cycle to cycle (*1–3*). Springs may also constrain animal performance, by making some frequencies of movement less energetically favorable than others. Any system with inertia and elasticity will have a resonant frequency of energy flow, defined by its mass, spring, and damping properties. An animal moving at its resonant frequency theoretically benefits from a larger kinematic output (i.e. limb motion) for the same actuation force input, since relatively small mechanical energy inputs each cycle compound to create a larger amplitude oscillation than would be possible at a non-resonant frequency (*4–6*). Conversely, operation at resonance may be detrimental for maneuverability, since an animal seeking to modulate its limb movement must work against a large amount of limb mechanical energy built up over many oscillations (*7*). The potential advantages and disadvantages of operating at resonance suggest that animals may balance trade offs in diversification of actuation and biomechanics across species with different behavioral and energetic requirements.

Resonance tuning may facilitate higher insect wingbeat frequencies, because mass-specific cost increases as size decreases for flapping organisms (*8–10*). Most flapping insects actuate their wings indirectly, through muscles that attach to the outside of a thin elastic exoskeletal shell (*11, 12*), which deforms to transmit muscle strain to rotational movement of inertial wings via a transmission known as the wing hinge. Flying insects move through the air, meaning that aerodynamically useful work done to support body weight is mostly dissipated to the surrounding fluid. As such, flying insects have been modeled as forced oscillators with elasticity, inertia, and nonlinear aerodynamic damping - a system with resonant mechanics (*5, 10, 13*). However, operating at resonance may also inhibit rapid frequency modulation of the wing stroke, which would require active muscular work against the movement of the wings and a temporary reduction of stroke amplitude. Frequency modulation may be important in agile flyers such as hawkmoths that rapidly maneuver in complex aerial environments (*7*). Evaluating this tradeoff broadly across insects remains an important challenge due to the lack of comparative measurements of spring-wing mechanics under a consistent model of resonance.

The aerodynamic efficiency-agility tradeoff implied by resonant wingbeats was pioneered by the work of Weis-Fogh in the mid 20th century, who defined a non-dimensional number (which we refer to as the Weis-Fogh number, *N*), measuring the ratio of peak inertial to aerodynamic torque over a wingstroke (*9, 13*). *N* has a convenient interpretation as the “sharpness” of an insect’s resonant curve(wingbeat amplitude vs. wingbeat frequency), analogous to the quality factor (*Q*) in engineered systems. Higher *N* implies a larger inertial power requirement relative to aerodynamics, and therefore both a larger potential benefit from elastic energy savings and steeper energetic penalty for off-resonance wingbeats. Lower *N* implies a larger relative aerodynamic power requirement and a larger fraction of total work done being used to support body weight, but a shallower resonant curve. The Weis-Fogh number provides a convenient nondimensional value for comparing resonant tradeoffs across species.

While there is little question that insects exhibit resonant mechanics (*8, 9, 14, 15*), direct measurements of resonant wingbeats have been limited to phylogenetically isolated species with disparate methods that complicate across-species comparisons (*5,16–19*). Furthermore, without independent, comparative measurements of each flight apparatus component, one cannot distinguish the specific traits that facilitate resonance tuning from traits that counteract resonance tuning due to competing constraints. For example, smaller (faster-flapping) insects actuate large amplitude wingstrokes with smaller muscle displacements, (*20*), and increasing their transmission ratio compared to larger insects (ratio of stroke angle amplitude to muscle displacement amplitude). As we demonstrate later, this in isolation would dramatically lower their resonant frequency (*5*), contrary to the expectation of resonance tuning. In general, it remains unknown how flight apparatus components scale with wingbeat frequency in closely related species and whether this variation contributes to evolutionary tuning of resonant mechanics.

The superfamily of moths Bombycoidea offers a unique opportunity to comparatively study the biomechanical drivers of insect resonant mechanics. Bombycoid moths represent over 5000 species and exhibit wide diversity in flight styles, feeding habits, body sizes, and wing morphology while maintaining similar component parts and actuation strategies (*21, 22*). Hawkmoths (family: Sphingidae) have evolved higher frequency wing beats and smaller, lower aspect ratio wings, diverging to a distinct adaptive peak from silkmoths. Many hawkmoths perform impressively agile hover-feeding behaviors, using their long proboscides to consume nectar while matching the flower’s position mid-air (*23*). Wild silkmoths (family: Saturniidae) have generally larger amplitude, lower frequency wing beats with larger wing areas and lack functional mouth parts as adults. Some also possess characteristic colour patterns and long wing tails and have evolved a more erratic flight style to evade predators (*24–26*).

Using Bombycoids as a model clade, we set out to answer two related questions:

1. Do individual components of the flight system relevant to resonance (i.e. stiffness, wing hinge transmission, wing inertia) scale with wingbeat frequency in accordance with resonant tuning? The resonant frequency of a spring-mass-damper typically increases with the square root of system stiffness and decreases with the inverse of the square root of inertia. Therefore, we predict these relationships for stiffness and inertia in moths, in agreement with resonance tuning. In particular, we predict the effects of stiffness and inertia to be strong enough to compensate for the higher wing hinge transmission ratio in smaller insects (*20*) that would in isolation reduce resonant frequency with higher wingbeat frequency. Taken together, we hypothesize that variation in these individual components combine to result in a resonant frequency that matches wingbeat frequency in Bombycoidea to maintain generally favorable resonant mechanics regardless of moth wingbeat frequency.
2. Do the resonant mechanics of hawk- and silkmoths reflect their different behavioral and energetic requirements? Given the stark clade-specific differences in wing morphology, kinematics, and feeding habits, we expect silkmoths to have more aerodynamically efficient resonant properties than hawkmoths, reflecting their frequently nutrient-limited adult life stage. We predict silkmoths to have a lower *N* than hawkmoths, resulting in a shallower resonance curve and providing them with a buffer of favorable frequencies around resonance at which to flap. As an alternative, silkmoths’ low *N* may preclude any meaningful energetic benefit to operating at resonance. Conversely, hawkmoths’ agile hover-feeding behavior leads us to predict they will be less constrained by precise matching of resonant and wingbeat frequencies, and have a larger *N*. Deviation from resonance may enable frequency-modulation at the expense of favorable energetics from perfectly resonant wingbeats. A larger *N* may allow them to return relatively more elastic energy than would be possible with a shallower resonance curve, even if operating off of resonance.

## 2 Materials and Methods

### Spring-wing resonance modeling framework

We build upon a recent single degree-of-freedom, lumped-parameter dynamics model of a flapping insect (*5, 13*). Newton’s second law for a rotational system with an aerodynamic force that is proportional to the magnitude of velocity squared (*5, 9, 27*) is:

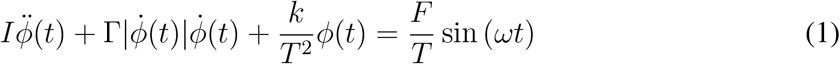

Here, *ϕ*(*t*) is the dynamic variable and represents the time-varying wingstroke angle. This single equation is parameterized by the linear thorax stiffness *k*, transmission ratio *T*, inertia of wings and added mass *I*, aerodynamic damping coefficient Γ, wing beat frequency *ω*, and muscle forcing amplitude *F*. The absolute value in the damping term ensures that the direction of the damping force always opposes wing motion. Note that the elastic term has the coefficient *k/T* ^2^, which we refer to as the wing hinge rotational stiffness *k*_*rot*_. We also assume both *k* and *T* are independent of wing angle, which we justify in the following sections. Since this equation is a non-linear second-order differential equation, we can numerically integrate to solve it for the wing stroke angle *ϕ*(*t*). Doing so over a range of potential frequencies *ω* yields a resonance curve (*ϕ* vs *ω*), the maximum of which is the displacement (damped) resonant frequency *f*_*res*_. We also are concerned with the undamped resonant frequency, sometimes called the natural frequency *f*_*nat*_, which has the convenient closed form:

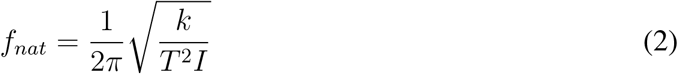

For lightly damped systems, *f*_*res*_ and *f*_*nat*_ are very close to one another, so their distinction is not important. However, since flapping insects are heavily damped by surrounding air, we compute both of these resonance frequencies which have slightly different physical meanings. We detail the measurement procedure or computation of each parameter in the following sections.

#### Animals

Live specimens from 10 species of Bombycoid moths were used in this study. Hawk-moth species used were *Manduca sexta, Smerinthus cerisyi, Hyles lineata, Hemaris diffinis, Sphinx chersis*. Silkmoth species used were *Actias luna, Automeris io, Antheraea polyphemus, Hyalophora cecropia, Citheronia regalis*. Since thorax material testing requires the animal be deceased, attempts were made to record muscle strain data from live animals before thorax stiffness experiments.

### Thorax stiffness measurements

We roughly followed the methods of precedent work with minor adjustments (*5, 16*). After anesthetizing in a refrigerator, moths were prepared for materials testing by removing their wings, legs, head, and abdomen. The third thoracic segment is not thought to be significantly responsible for wing actuation or elastic energy storage, and was removed to expose the posterior phragma. Waxy cuticle was gently filed off from the scutum and phragma to support better glue adhesion. Finally, the thoracic ganglion and thoracic musculature were severed to prevent spontaneous muscle contractions from transferring appreciable force to the force sensor.

A custom 3D printed mount was secured to the anterior muscle attachment surface on the scutum with cyanoacrylate glue. The scutum mount was rigidly attached to the drive shaft of an electrodynamic shaker (The Modal Shop 2007E). A #2 threaded rod was glued to the posterior phragma, and was screwed into the insert of a piezoelectric force transducer (PCB Piezotronics 209C11). The force transducer was attached to a micromanipulator, giving us precise control over the thorax’s rest length. We defined rest length as the length at which no force was measured by the transducer. Importantly, we aligned the thorax such that deformations occurred along the axis that the downstroke muscle would contract in the live animal (Fig. 1a). A fiber-optic displacement sensor (Philtec D47) was used to measure displacement from a strip of reflective tape attached to the shaker drive shaft. The displacement sensor was used only in its linear range, and was calibrated daily immediately prior to mounting the first specimen. We prescribed a sine chirp displacement signal from 1-100 Hz at 9% peak-to-peak strain amplitude (*28*). The same thorax strain amplitude was used for all species since species-specific strain data could not be analyzed prior to performing stiffness measurements. We do not consider contributions of active muscle stiffness which are relatively small in hawkmoths compared to exoskeletal stiffness (*5, 29*). Recent work on the thorax of *Manduca sexta* has demonstrated that stiffness and material damping in the thorax are frequency-independent (*12, 30*). As such, we ignore internal damping in resonance calculations, as it does not affect the location of the resonant peak, and we use the stiffness measured at wingbeat frequency as the thorax stiffness for each species. Our sign convention is such that positive force is generated in the shortening direction.

**Figure 1.**
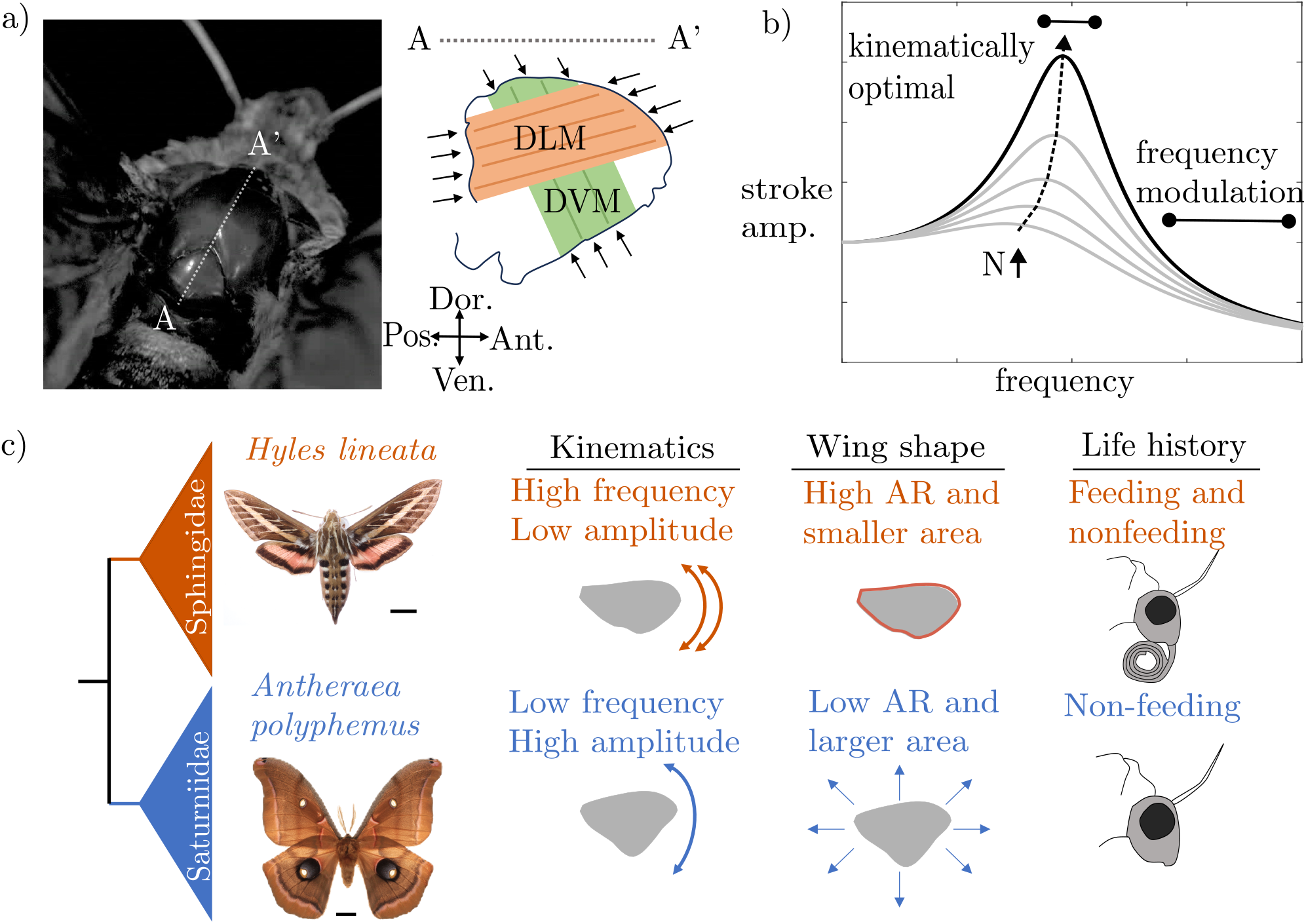
a). Photograph of a hawkmoth thorax with the cross-section along line AA’ shown to the right. The downstroke (DLM) and upstroke (DVM) muscles are shown, along with their lines of action. Modified with permission from Gau et al. 2022 (*5*) b). Generalized resonance curve for a spring-mass-damper system, showing regimes of kinematically optimal and more maneuverable wingbeat frequencies. Grey curves show the effect of increasing *N*. c). Hawk-moths (Sphingids) and silkmoths (Saturniids) are sister families that make up the superfamily Bombycoidea. The two clades exhibit different flight kinematics, wing morphologies, and life history traits. Modified with permission from Aiello et al. 2021 (*21*)

### Muscle strain measurements

Moths were anaesthetized in a refrigerator, tethered ventrally, and positioned beneath a high-speed video camera (Photron FastCam mini UX100). The moth’s abdomen and third thoracic segment were removed, exposing the posterior phragma, the attachment point for the main downstroke muscles (DLMs). Scales were removed from the anterior scutum, and a white paint pen was used to mark muscle attachment points on each side of the animal. The moth was stimulated to flap by gentle tactile stimulation on the back of its neck. After a flapping bout, the camera was moved to capture wingbeats from the front of the animal, so that wingbeat amplitude could be measured. The order of recording head-on and top-down angles was randomized from individual to individual to ensure moths were not excessively fatigued during a particular filming angle. White markers and wing tips were digitized in DLTdV8 over multiple wingbeats (*31*). We define a sign convention such that positive strain is shortening.

In general, we found that tethered moths flapped with wingbeats that were larger in amplitude than those in free-flight (*21*), likely because of the highly invasive procedure necessary to expose the posterior phragma and stress due to tethering. To ensure that strain and transmission calculations were not affected by these inflated amplitudes, we computed transmission ratio (T) directly from the phragma displacement *d*_*max*_ and front-view wingstroke amplitude *ϕ*_*tethered*_ measurements in each moth individually.

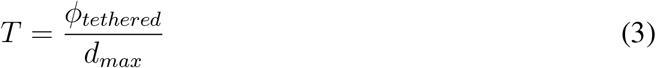

Computing transmission ratio in this way using maximum wingbeat amplitude and maximum strain from the same individual ensure that any inflated wingbeat amplitude is matched by an inflated thorax strain. For a linear and frequency-independent transmission, this will result in an accurate estimation of the transmission ratio. In *Manduca*, we confirmed that the transmission of the wing hinge is linear by recording wingbeats with a time-synchronized two-camera setup (Fig. 2d). Plotting wing angle as a function of thorax displacement results in an ellipse, indicating that a linear transmission is likely an appropriate assumption for this group of animals. We defined the operating length *L*_*op*_ of the thorax as the mean strain across all digitized wingbeats. We then used *T*, *L*_*op*_, and free-flight wingbeat amplitude *ϕ*_*o*_ to calculate a muscle strain likely to be generated in free-flight by the following equation:

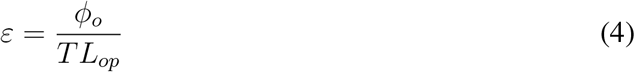

Results of our method for *Manduca* result in a transmission ratio in rough agreement previous calculations using data from a different study of muscle length changes in a tethered animal (*5, 28*). For analysis of strain and transmission data, each flight bout from an individual was considered a separate trial. Data was collected from at least three different individuals of each species and was pooled for species-averaged analysis.

**Figure 2.**
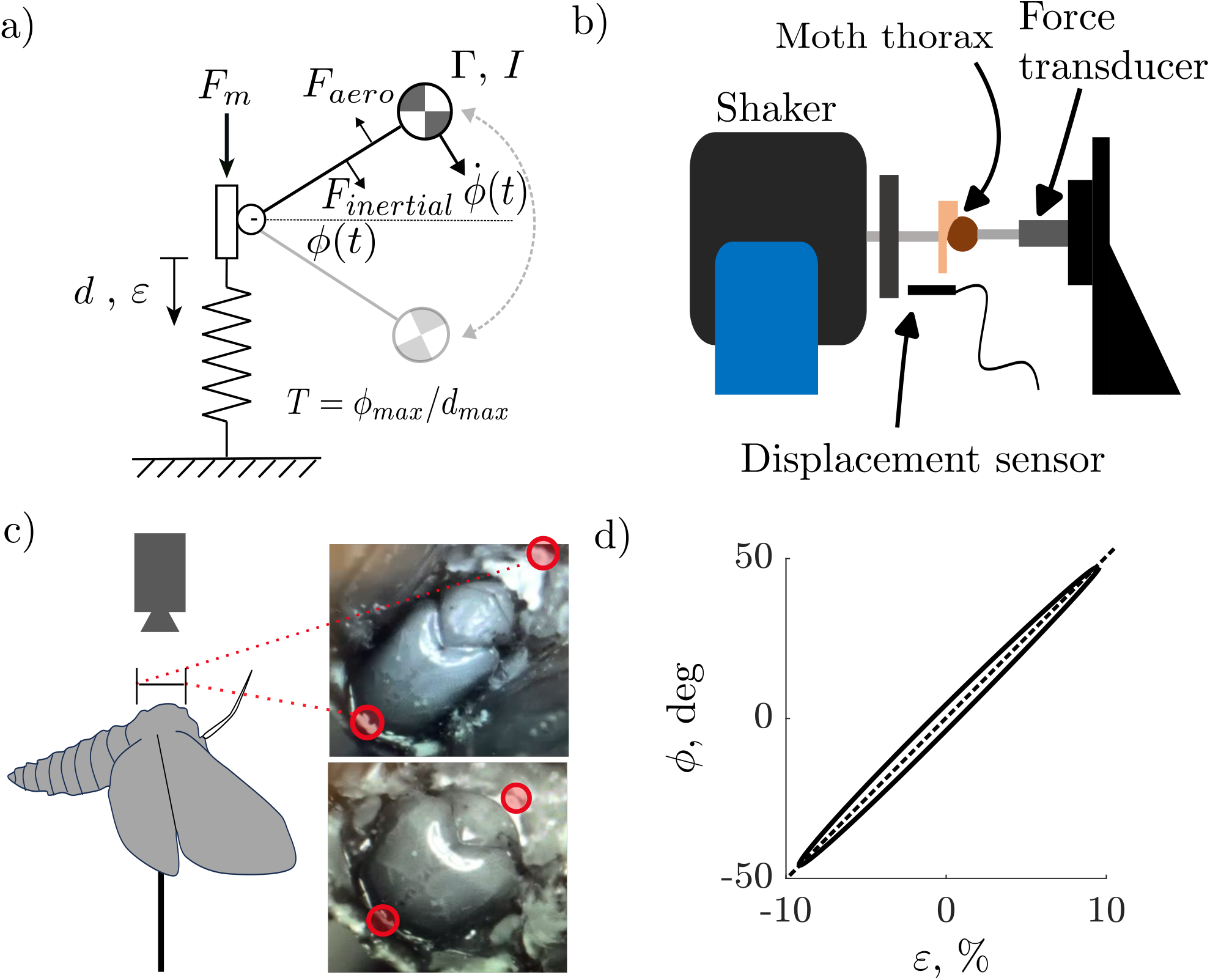
a). Free-body diagram of generalized spring-wing model showing the geometry of the actuating muscles, thoracic spring, transmission, and inertial wing. b). Schematic of the experimental apparatus used to measure thorax stiffness, consisting of an electrodynamic shaker, fiber-optic displacement sensor, and piezoelectric force transducer. c). Schematic of a moth filmed from above on a tether, with inset photographs showing the DLM attachment points at maximum and minimum strain. d). Average wingstroke angle as a function of thorax strain in one representative *Manduca* individual. The transmission ratio (slope of the ellipse denoted by dashed line) does not deviate substantially from linearity over a wingstroke.

### Inertia and aerodynamic damping calculation

We leveraged an existing wing morphometric dataset to calculate inertial and aerodynamic parameters for each moth species (*22*). Specific imaging and digitization methods are identical to those in Aiello et al. 2021. For *Hemaris diffinis, Smerinthus cerisyi, and Sphinx chersis*, we utilized wing shape data from a similarly sized and closely related species in the same genus: *Hemaris thetis and Hemaris thysbe, Smerinthus ophthalmica and Smerinthus jamaicensis, and Sphinx kalmiae*. In the above cases, distinctions in wing morphology between species in the same genus were minimal and should not meaningfully affect our results. We compute inertia of the wing pair and added air mass from the following equation (*32*), where *m*_*w*_ is the mass of the wing pair, *R* is wing length, 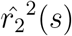 is the second moment of wing shape, *ν* is the wing added mass, and 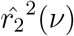 is the second moment of added mass (see Table 2 for a complete glossary of symbols):

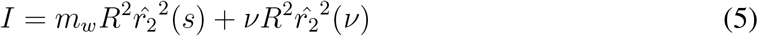

*ν* was computed from the following equation, where 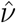 is the non-dimensional added mass and *Ar* is the aspect ratio:

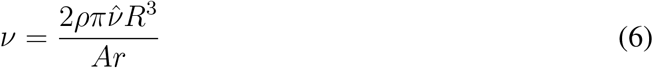

With this measure of inertia, we computed the fraction of inertial work offset by elastic energy storage. Integrating the elastic and inertial terms of eq. (1) over a quarter-stroke yields the work contributions 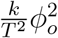 and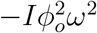. We considered only a quarter-stroke since the symmetry of a sinusoidal wingstroke means that the calculation will be the same. Taking the quotient of these expressions gives us an estimate of the amount of energy returned by the spring relative to its potential maximum benefit.

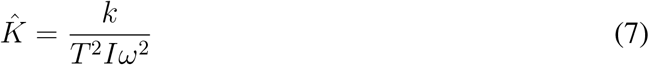

We utilized a constant quasi-static aerodynamic damping model with velocity-squared damping, such that the aerodynamic force over a wingstroke is equal to 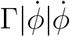. This simplified representation of flapping-wing aerodynamics has been used previously to model moth resonant mechanics and its functional form allows for derivation of analytical results that would be impossible with more complex models (*5,9,27,33*). In later sections, we discuss how more realistic aerodynamics may be considered. The aerodynamic damping parameter Γ is computed by the following equation (*27*), where *ρ* is the air density, *C*_*D*_ is the average wing drag coefficient, *A*_*w*_ is the area of the wing pair, 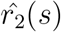is the non-dimensional second-moment of wing shape, *R* is wing length, and *l*_*cp*_ is the non-dimensional location of the center of pressure.

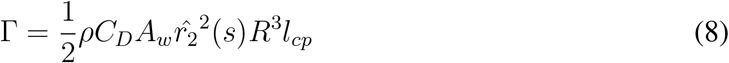

*C*_*D*_ was computed for each species using the equation 3.2 of Han et al. 2015, averaged over the wingstroke using angle-of-attack data from free-flight kinematics for each species (see following section) (*34*). A constant *l*_*cp*_ of 0.6 was used for all species (*27*).

**Table 1:**
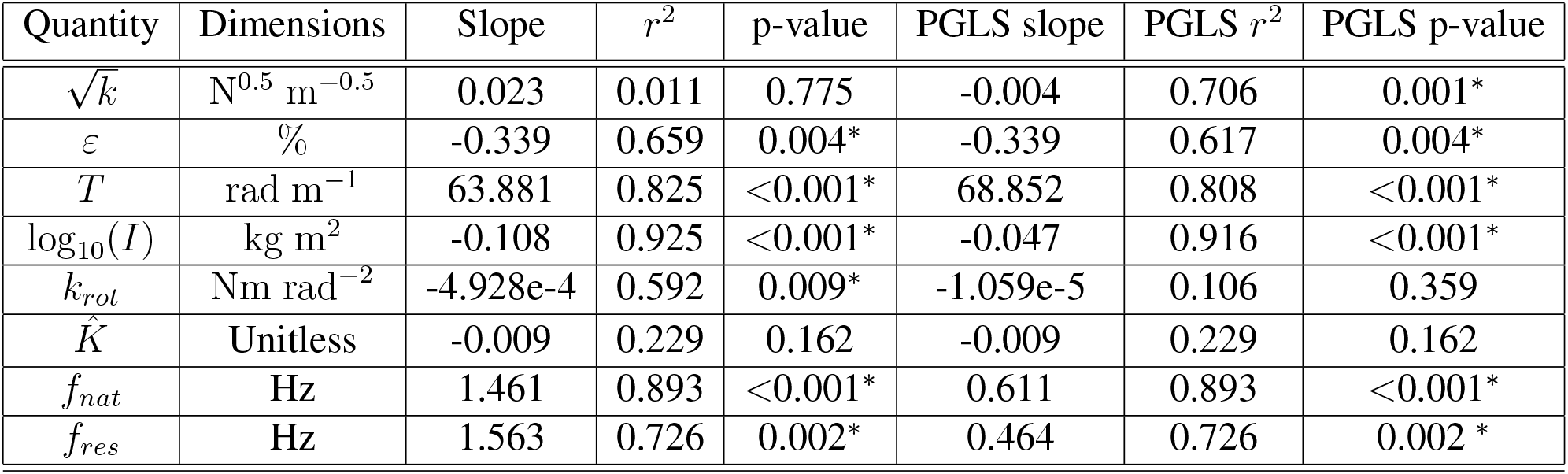
Regression and PGLS slopes and *R*^2^ values for each quantity in Figs 3–5 plotted against wingbeat frequency.

| Quantity | Dimensions | Slope | $r^2$ | p-value | PGLS slope | PGLS $r^2$ | PGLS p-value |
| --- | --- | --- | --- | --- | --- | --- | --- |
| $\sqrt{k}$ | $\text{N}^{0.5} \text{m}^{-0.5}$ | 0.023 | 0.011 | 0.775 | -0.004 | 0.706 | 0.001* |
| $\varepsilon$ | % | -0.339 | 0.659 | 0.004* | -0.339 | 0.617 | 0.004* |
| $T$ | $\text{rad m}^{-1}$ | 63.881 | 0.825 | <0.001* | 68.852 | 0.808 | <0.001* |
| $\log_{10}(I)$ | $\text{kg m}^2$ | -0.108 | 0.925 | <0.001* | -0.047 | 0.916 | <0.001* |
| $k_{rot}$ | $\text{Nm rad}^{-2}$ | -4.928e-4 | 0.592 | 0.009* | -1.059e-5 | 0.106 | 0.359 |
| $\hat{K}$ | Unitless | -0.009 | 0.229 | 0.162 | -0.009 | 0.229 | 0.162 |
| $f_{nat}$ | Hz | 1.461 | 0.893 | <0.001* | 0.611 | 0.893 | <0.001* |
| $f_{res}$ | Hz | 1.563 | 0.726 | 0.002* | 0.464 | 0.726 | 0.002* |

**Table 2:**
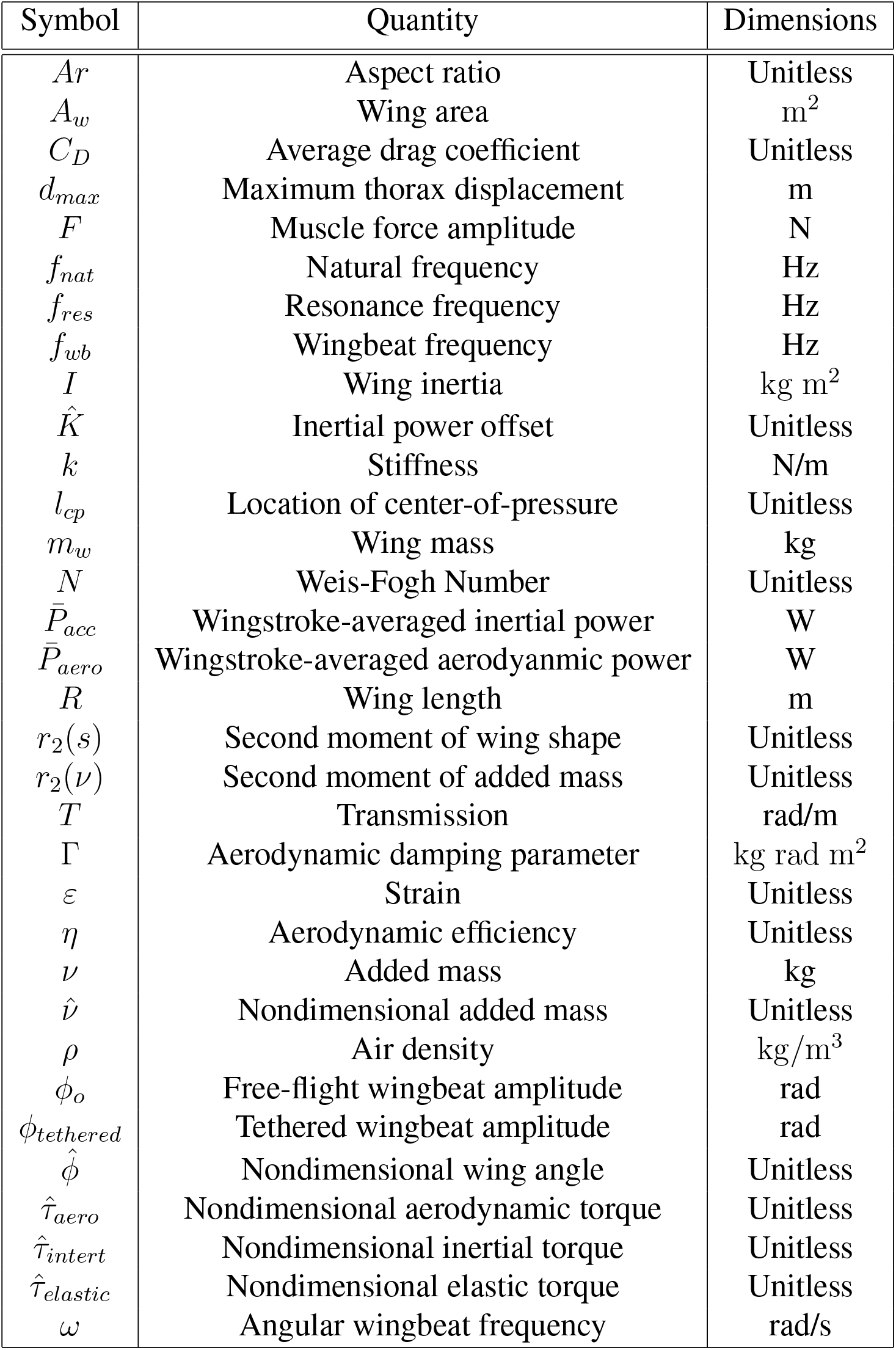
Symbol glossary.

| Symbol | Quantity | Dimensions |
| --- | --- | --- |
| $Ar$ | Aspect ratio | Unitless |
| $A_w$ | Wing area | $m^2$ |
| $C_D$ | Average drag coefficient | Unitless |
| $d_{max}$ | Maximum thorax displacement | m |
| $F$ | Muscle force amplitude | N |
| $f_{nat}$ | Natural frequency | Hz |
| $f_{res}$ | Resonance frequency | Hz |
| $f_{wb}$ | Wingbeat frequency | Hz |
| $I$ | Wing inertia | $kg\ m^2$ |
| $\hat{K}$ | Inertial power offset | Unitless |
| $k$ | Stiffness | N/m |
| $l_{cp}$ | Location of center-of-pressure | Unitless |
| $m_w$ | Wing mass | kg |
| $N$ | Weis-Fogh Number | Unitless |
| $\bar{P}_{acc}$ | Wingstroke-averaged inertial power | W |
| $\bar{P}_{aero}$ | Wingstroke-averaged aerodynamic power | W |
| $R$ | Wing length | m |
| $r_2(s)$ | Second moment of wing shape | Unitless |
| $r_2(\nu)$ | Second moment of added mass | Unitless |
| $T$ | Transmission | rad/m |
| $\Gamma$ | Aerodynamic damping parameter | $kg\ rad\ m^2$ |
| $\varepsilon$ | Strain | Unitless |
| $\eta$ | Aerodynamic efficiency | Unitless |
| $\nu$ | Added mass | kg |
| $\hat{\nu}$ | Nondimensional added mass | Unitless |
| $\rho$ | Air density | $kg/m^3$ |
| $\phi_o$ | Free-flight wingbeat amplitude | rad |
| $\phi_{tethered}$ | Tethered wingbeat amplitude | rad |
| $\hat{\phi}$ | Nondimensional wing angle | Unitless |
| $\hat{\tau}_{aero}$ | Nondimensional aerodynamic torque | Unitless |
| $\hat{\tau}_{inert}$ | Nondimensional inertial torque | Unitless |
| $\hat{\tau}_{elastic}$ | Nondimensional elastic torque | Unitless |
| $\omega$ | Angular wingbeat frequency | rad/s |

### Moth free-flight kinematics

We leveraged existing wind tunnel free-flight videos of moths, which were collected and digitized as described by Aiello et al. 2021 (*21*). Wingstrokes from multiple individuals were averaged to yield a species-specific frequency and stroke plane sweep-angle amplitude. For *Sphinx chersis, Hemaris diffinis, and Smerinthus cerisyi*, we utilized free-flight data from similarly sized and closely related species of the same genus *Sphinx kalmiae, Hemaris thysbe, and Smerinthus ophthalmica* instead. Time-varying angle-of-attack data for each species was also used to compute a wingstroke-averaged species-specific drag coefficient as outlined in the previous section.

### Simulation

Using the parameters from each species, we simulated a frequency sweep from 1-100 Hz using species-averaged parameter values for *k, T*, *I*, and Γ. Muscle forcing amplitude *F* was computed for each species by selecting the value that results in a wing beat amplitude that matches free-flight experiments for a simulated insect driven at its free-flight wing beat frequency. This method accounts for the fact that in vitro muscle physiology experiments on moths have resulted in force estimates that are over an order of magnitude lower than what would be required to sustain flight (*5, 29, 35*). Resonant frequency (*f*_*res*_) is computed as the frequency at which maximum steady-state wingbeat amplitude occurs.

### The Weis-Fogh number and aerodynamic efficiency

The Weis-Fogh number was originally defined as the ratio of peak inertial to peak aerodynamic torque over a wingstroke, given by the following equation:

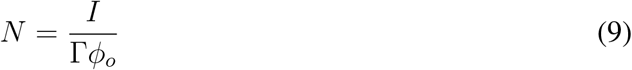

Using this equation to compute *N* across species relies upon a heavily simplified aerodynamic model with a constant damping parameter. As a way to incorporate slightly more aerodynamic realism we computed *N* using wingstroke-averaged aerodynamic and inertial power computations from a recent blade-element model (*21*). See supplementary information for a derivation of this equation.

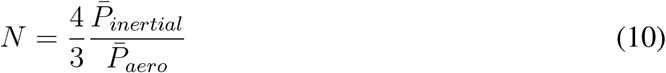

Weis-Fogh originally defined aerodynamic efficiency of a flapping insect as the fraction of total work required for flight taken up by aerodynamic costs (*9*). Aerodynamic costs represent the ‘useful’ work that contributes to supporting body weight, while inertial and elastic costs are necessary to actuate flight but do not aid in body weight support. We introduce two non-dimensionalizations to make the computation of these energy costs easier (*13*). First, we non-dimensionalized the wing angle in equation (1) by the wingstroke amplitude such that 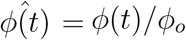. We then non-dimensionalized each torque in equation (1) by the peak aerodynamic torque at midstroke, resulting in the following non-dimensional torques as a function of non-dimensional wing angle 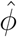:

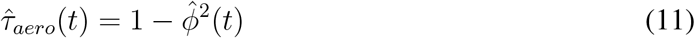

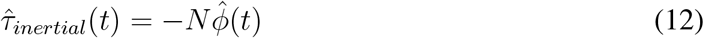

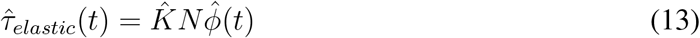

We can then integrate each torque to yield the work associated with each energy cost. We only consider positive work, as negative work done by muscles incurs significantly less metabolic energy cost and is often modeled as negligible (*10, 36*). Choosing the bounds of integration such that only positive work is considered (Supplementary S1), we can compute aerodynamic efficiency by the following equation:

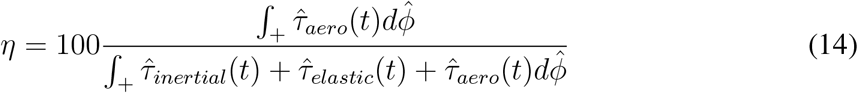

This definition of aerodynamic efficiency is distinct from any notion of metabolic efficiency, which we are not directly assessing in the current work.

### Statistics

We use two-sample t-tests to compare clade-dependent differences (grouped as hawkmoths and silkmoths) in resonant mechanical properties with a significance threshold of 0.05. For continuous data, we perform linear regressions and illustrate a line and confidence intervals only if we find a significant relationship. In addition, we perform Phylogenetic Least Squares (PGLS) to confirm that any apparent trend is cannot be attributed to phylogenetic distance between species alone. We implement PGLS in R with the phytools and caper packages.

## 3 Results

### 3.1 Thorax stiffness does not increase with wingbeat frequency

To identify how components of the flight system contribute to resonant mechanics across moths of varying wingbeat frequency, we measured stiffness, transmission, and inertia comparatively in hawk- and silkmoths. We performed dynamic material testing over a range of frequencies to measure thorax stiffness, extracting the stiffness of the thorax at wingbeat frequency for each moth (Fig. 3a). We found that stiffness does not vary significantly across the nearly order of magnitude of wingbeat frequency variation captured in our species sampling, and there are no detectable differences between hawkmoths and silkmoths when grouped by clade (Fig. 3b). Substantial inter-subject variation is persistent across species (Fig 3a). Since the resonant frequency of a spring-mass-damper is proportional to the square root of stiffness, we expected 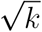 to scale with the wingbeat frequency of an insect flapping at resonance. Howeve 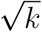, we could not detect a statistically significant relationship between wingbeat frequency and 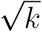. When controlling for phylogenetic relatedness using Phylogenetic Generalized Least Squares (PGLS), we find a significant relationship but with a slope that suggests negligible biological significance (predicts less than a percent decrease in *k* between fastest and slowest insects measured) (Fig 3c, Table 1). Stiffness measurements were lower than some previous measurements, which we attribute to the fabrication of 3D printed mounts that better match the DLM attachment surfaces, thus resulting in more realistic deformations (*12, 30*). The current results are in close agreement with recent independent measurements from *Manduca* (*37*). Regardless, stiffness does not co-evolve with wingbeat frequency.

**Figure 3.**
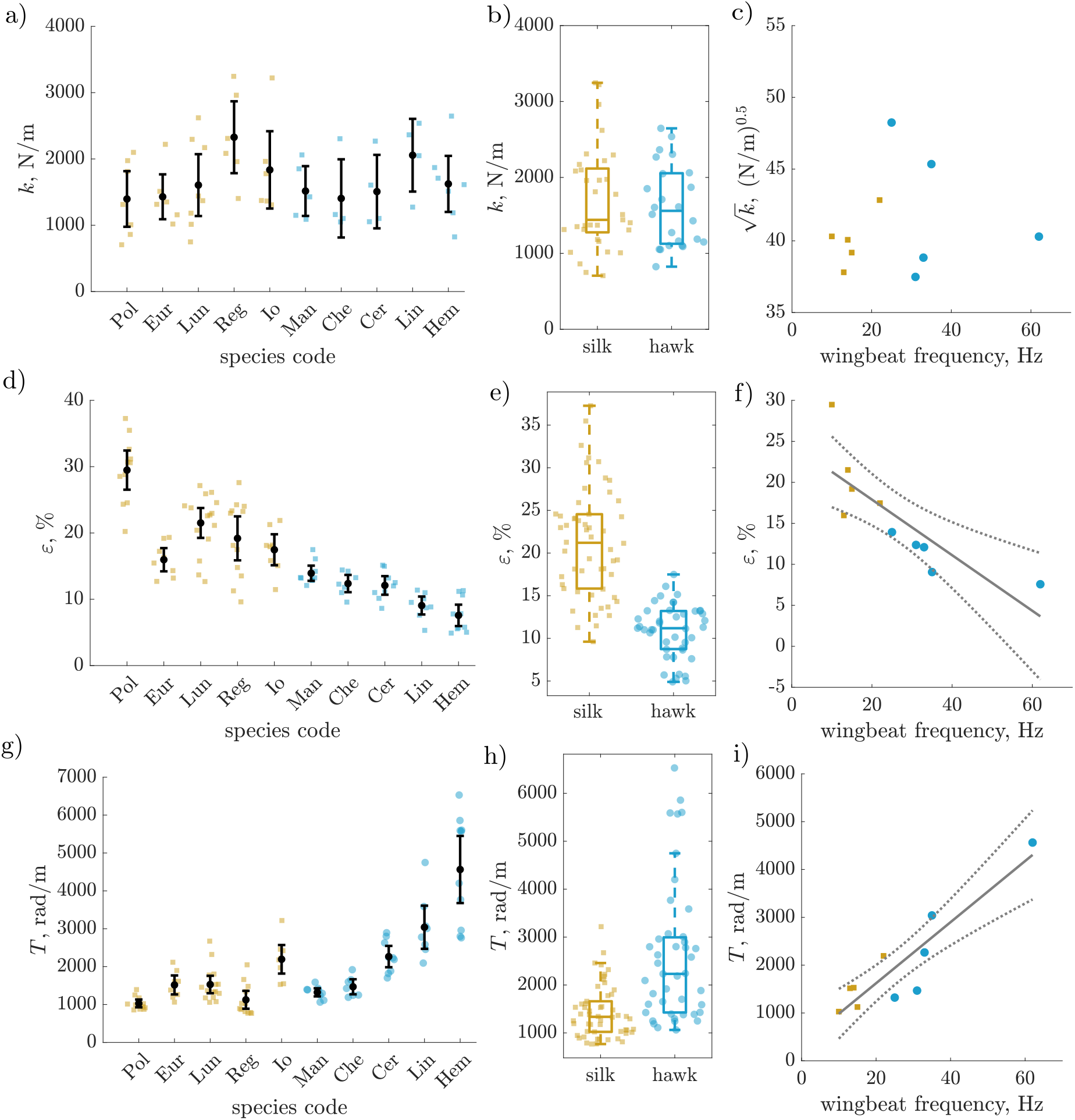
a). Stiffness does not show a distinct pattern of variation across species. Each marker represents a different individual, with orange square corresponding to silkmoths and blue circles corresponding to hawkmoths. Black circles are species means and error bars show 95% confidence intervals of the mean. b). Stiffness does not differ significantly between clades within Bombycoidea. c). There is no significant relationship between 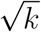 and wingbeat frequency. d). Thoracic strain varies between species. e). Silkmoths have higher average strains than hawk-moths. f). Strain varies with wingbeat frequency in an inverse fashion. g). Transmission ratio varies strongly with species. h). Hawkmoths have higher transmission ratios than silkmoths. i). Transmission ratio varies in a positive linear fashion with wingbeat frequency

### 3.2 Muscle strain decreases and transmission ratio increases with wing-beat frequency

We next examined whether muscle strain and transmission ratio vary with wingbeat frequency in a way that facilitates resonance tuning. Muscle strain amplitude varied inversely with wing-beat frequency, with over 20% difference between lowest and highest frequency animals (Fig. 3d,f). *Antheraea polyphemus* exhibited the largest muscle strain amplitude, with individuals exceeding 30%, a large peak-to-peak strain for a cyclic movement at around 10 Hz. Other slow-flapping silkmoths we measured only exhibited strains of 16 *−* 25% (Fig. 3d), suggesting *A. polyphemus* has muscles capable of particularly high strain. On average, silkmoths exhibited over two-fold larger strains than hawkmoths (Fig 3e). Transmission ratio is a composite quantity that takes muscle strain, thorax length, and wingbeat amplitude into account. We find transmission scales roughly linearly with wingbeat frequency (Table 1), such that higher frequency moths have a larger transmission ratio (Fig 3g-i). This follows directly from the muscle strain result because muscle strain and transmission ratio are inversely proportional. Between our fastest and slowest moths, transmission ratio varied by a factor of five (Fig. 3i).

### 3.3 Hawkmoths and silkmoths both offset inertial power costs with elastic energy storage

Combining thorax stiffness and transmission ratio into the wing rotational stiffness (*k*_*rot*_), we find a weak inverse relationship between rotational stiffness and wingbeat frequency (Fig. 4b) that is not supported under PGLS (Fig 4, Table 1). This result is contrary to common intuition that faster oscillators are stiffer, as well as some previous scaling predictions for rotational stiffness as a function of body size (*38*). Wing inertia falls off sharply and in a nonlinear fashion with wingbeat frequency, ranging over 2.5 orders of magnitude (Fig. 4a). This decrease is much steeper than what would be expected from the *−* 0.5 exponent predicted by Eq. 2. From Eq. 5, we compute the fraction of inertial power offset by elastic energy storage from rotational stiffness and wing inertia. Surprisingly, these two quantities combine to result in no discernable relationship between inertial power offset and wingbeat frequency (Fig. 4c, Table 1). Both hawk and silkmoth thoraces return substantial energy from cycle-to-cycle, with the lowest 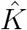 hawkmoth still offsetting 20% of its inertial power costs. The slowest moth we tested, *Antheraea polyphemus*, has a 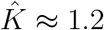, indicating that its thorax elasticity is slightly overtuned with respect to its inertial costs (Fig. 4c).

**Figure 4.**
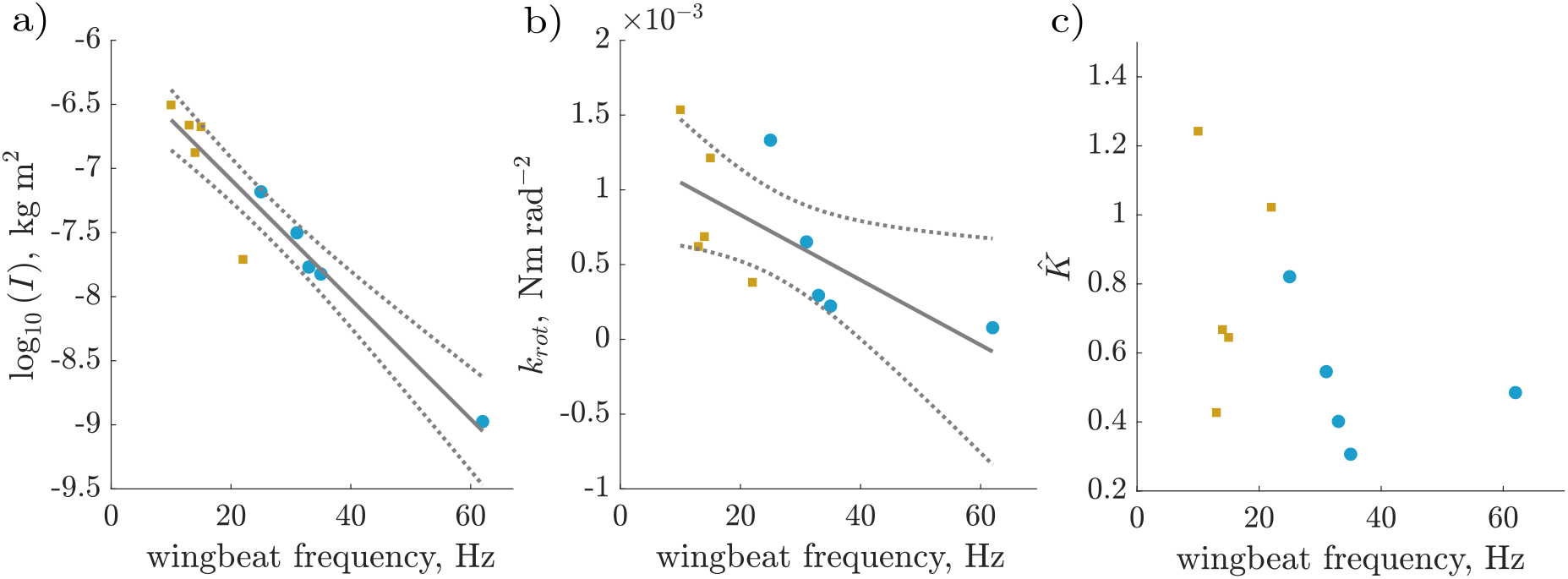
a). The logarithm of wing inertia decreases sharply with frequency (regression: *r*^2^ = 0.925, *p* =*<* 0.001^*∗*^; PGLS: *r*^2^ = 0.916, *p* =*<* 0.001^*∗*^). b). Linear regression of rotational stiffness vs wingbeat frequency shows a weak negative relationship (*r*^2^ = 0.592, *p* = 0.009^*∗*^), but this relationship becomes insignificant when controlling for phylogeny (PGLS *p* = 0.359). c). Moth species offset widely varying proportions of their inertial power costs with elastic energy storage and return, although on average the energetic benefit is substantial,approximately 66*±*30%. There is no significant relationship between wingbeat frequency and 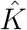 (PGLS *p* = 0.162). In all plots, linear regression lines are shown in solid grey, with 95% confidence intervals shown in dotted grey. All individual points correspond to species-average values.

### 3.4 Bombycoid resonant mechanics reflect behavioral and energetic requirements

Combining measurements of thorax properties, muscle strain, wing morphology, and free-flight kinematics, we can compute resonance frequencies for each species of moth from our spring-wing model (Eq. 1-2). There are multiple possible resonant frequencies. We find a correlation between moth wingbeat frequency and both undamped (velocity) (linear regression: *r*^2^ = 0.893, *p* =*<* 0.001^*∗*^; PGLS: *r*^2^ = 893, *p <* 0.001^*∗*^) and damped (displacement) (linear regression:*r*^2^ = 0.726, *p* = 0.002^*∗*^; PGLS: *r*^2^ = 0.726, *p* = 0.002^*∗*^) resonance frequencies (Fig. 5a). In general, we find that undamped resonance frequency is a good predictor of wingbeat frequency across species, as evidenced by most species lying very close to the equivalency line in Fig. 5a. Damped resonance frequencies lie farther from wing beat frequencies because consideration of aerodynamic damping reduces the system resonant frequency further. Regardless of which resonance frequency is being considered, hawkmoths lie at least 10 Hz farther from resonance on average than silkmoths (damped: *p* = 0.008, undamped: *p* = 0.0155) (Fig. 5b-c).

**Figure 5.**
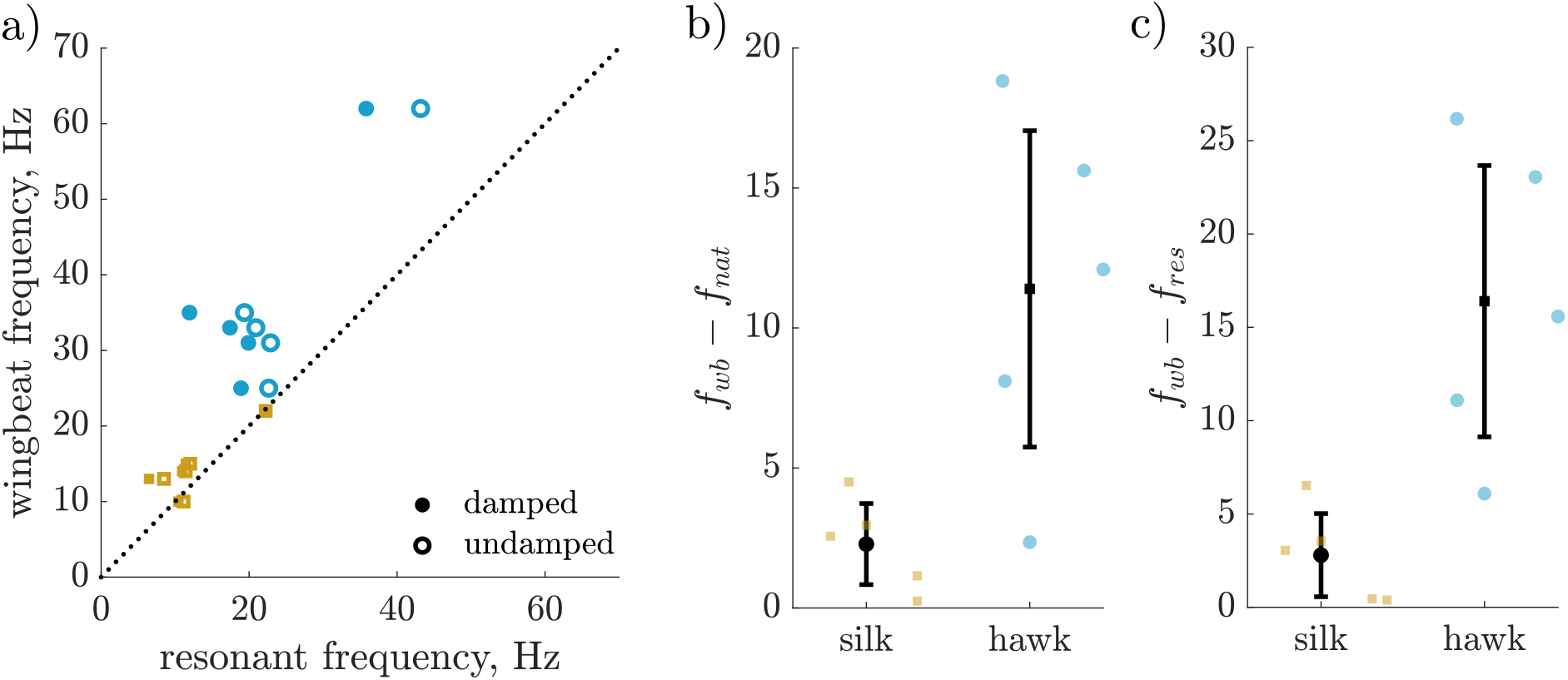
a). Both undamped and damped resonance frequencies of moths lie above the equivalency line, indicating that Bombycoid moths are generally supra-resonant. b-c). The dfference between wingbeat frequency and resonance frequency (undamped and damped) is larger in hawkmoths than silkmoths.

While operation close to a resonant peak is generally indicative of efficiency, resonant frequency alone is not enough to quantitatively evaluate a moth’s aerodynamic efficiency. This is because efficiency also depends on the shape of an insect’s resonance curve, which can be captured by the Weis-Fogh number (*N*). An insect with a wider resonance curve (lower *N*) will incur lesser relative energetic penalties for operating off of resonance. To evaluate the contributions of resonance curve shape and distance from the resonant peak to overall animal performance, we compute each moth species’ aerodynamic efficiency. Weis-Fogh originally defined this quantity as the ratio of aerodynamic work to total work required by an insect over a wingstroke (*9*). Using Eq. 1, we can write expressions for non-dimensional torques about the wing hinge as a function of non-dimensional wing angle. The energy cost associated with each torque is then represented by the integral of that torque with respect to wing angle. We assume that only positive work contributions require significant metabolic energy, and compute aerodynamic efficiency as the ratio of blue and grey areas in Fig. 6a-c (a more detailed mathematical description of this calculation is in the supplement). We diagrammatically show the integrals of Eqs. 10-12 in Fig. 6 in three resonant regimes: above, equal to, and below undamped resonance. Above and below resonance, inertial torque exceeds elastic torque during a portion of each halfstroke, resulting in a positive work cost (green shaded area and bars in Fig. 6a,d,c,f) that must be supplied by musculature. At resonance, inertial and elastic torque cancel exactly at all 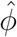 so that the only mechanical work required of the musculature is due to aerodynamics (Fig. 6b,e).

**Figure 6.**
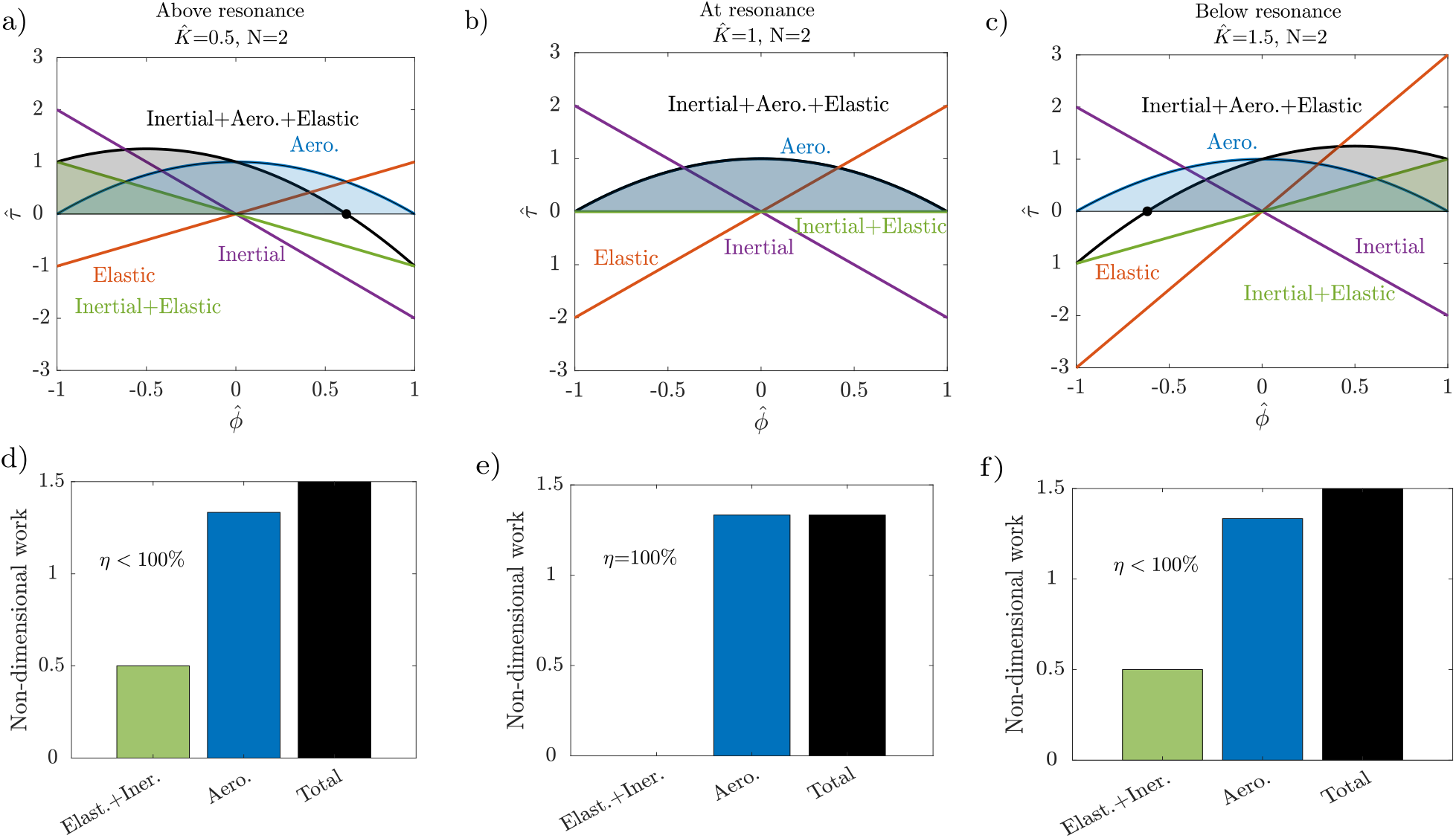
Non-dimensional torques and work at three different resonant conditions. a-b) Above (faster than) resonance, the magnitude of inertial torque exceeds elastic torque, requiring net positive mechanical work during the first half of each halfstroke (green shaded area and bar). As such, aerodynamic efficiency is below 100%. c-d) At resonance, inertial and elastic torques cancel exactly at every point during the wingstroke, so the only source of net positive work is due to aerodynamics. As such, aerodynamic efficiency is 100%. e-f). Below resonance, elastic torques exceed inertial torques requiring net positive work during the second half of each halfstroke (green shaded area and bar). This results in the same efficiency as in the above resonance case.

Having established a measure of aerodynamic efficiency that depends on *N*, we use Eq. 10 to compute *N* comparatively. We estimate that *N* values for our moths lie between 1 and 5, in agreement with previous estimates for similar insects (Fig. 7a). Importantly, each species had 1 *< N <* 5, indicating potential for elastic energy offset of inertial costs, but with a relatively shallow resonance curve, especially when compared to engineered oscillators which have quality factors that far exceed 10 (*39*). We find that hawkmoths have a larger *N* on average than silkmoths (Fig. 7a). When we calculate aerodynamic efficiency using Eq. 13, the resulting efficiency space depends only on two parameters, *N* and 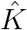 A maximum occurs at 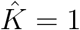 which corresponds to operation at undamped resonance (Fig. 7d). Efficiency decreases with increasing *N*, with this effect being steeper farther away from resonance. A smaller *N* implies a larger relative aerodynamic cost, thus a greater fraction of the insect’s per-wingstroke energy budget is being devoted to useful aerodynamic work. When we place hawkmoths and silkmoths on this efficiency space, we find that they cluster based on their differing 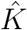 and *N* values. We find silkmoths have 10% higher aerodynamic efficiency on average than hawkmoths (*p* = 0.0190), mostly due to their lower average *N* (Fig. 7c) (*p* = 0.0033). Hawkmoths’ higher *N* places them in a location in the space with a steep efficiency gradient. Therefore, their supra-resonant wingbeats incur a larger efficiency penalty than would be the case at a lower *N*.

**Figure 7.**
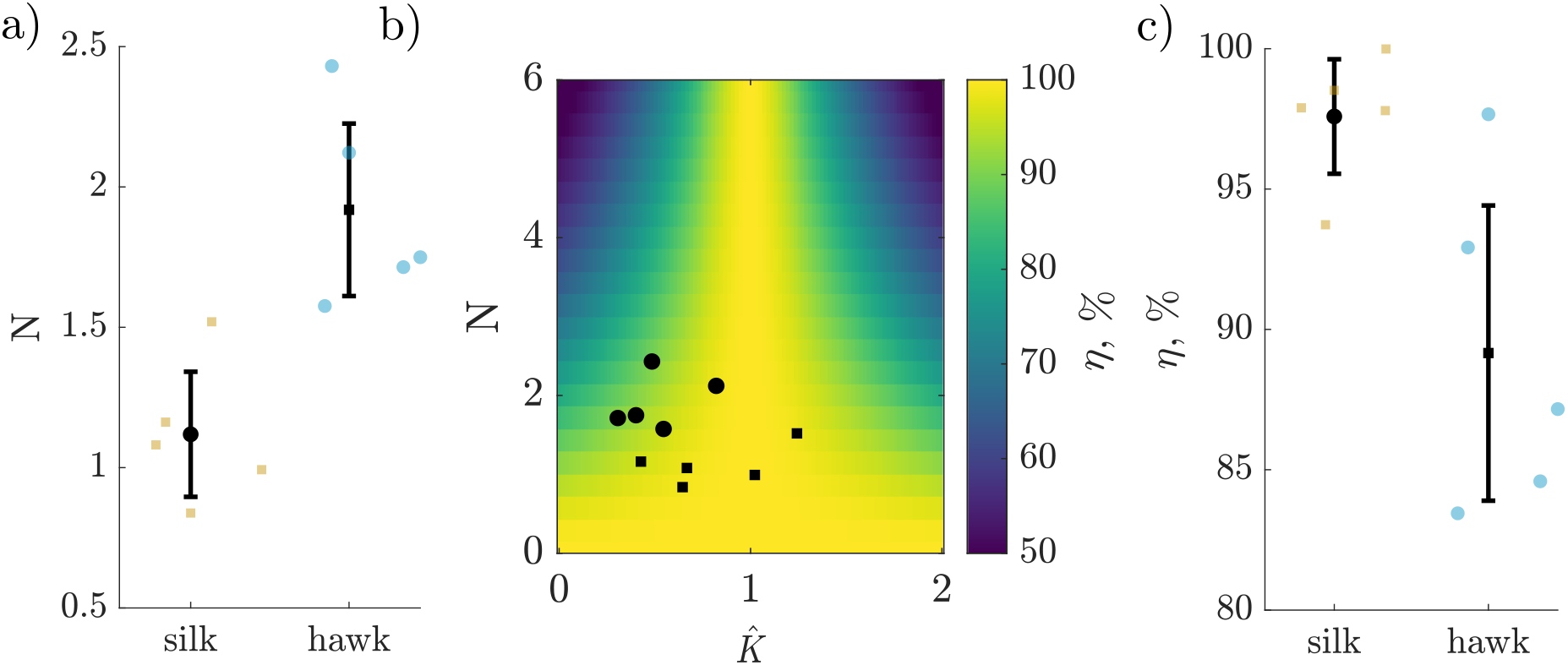
a). Weis-Fogh number (*N*) is larger in hawkmoths than silkmoths (*p* = 0.0033). b). Aerodynamic efficiency space for resonant flappers depends on *N* and 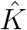. Hawkmoths (large dots) and silkmoths (small squares) cluster in this space. c). Combining resonant frequency and the Weis-Fogh number for each moth species reveals that silkmoths have higher aerodynamic efficiencies on average than hawkmoths (*p* = 0.0190).

## 4 Discussion

Our first objective in this study was to understand whether components of the flight system vary with wingbeat frequency in a way that facilitates or works against resonance tuning in moths. We find strong support for our hypothesis that wingbeat frequency scales with resonant frequency, but not because of the expected scaling of stiffness. Our results demonstrate no significant scaling relationship between thorax stiffness and wingbeat frequency (Fig. 3a- c), but a positive linear relationship between transmission and wingbeat frequency (Fig 3d-f). Combined, these trends result in a wing hinge rotational stiffness that decreases with wingbeat frequency (Fig. 4b). This trend alone is contrary expected positive scaling of rotational stiffness to result in resonant tuning. However, due to the sharply nonlinear inverse scaling of wing inertia with wingbeat frequency (Fig. 4a), resonance frequency scales linearly with wingbeat frequency (Fig 5a). Moreover, in most cases wingbeat frequencies tend to be offset such that they exceed the resonant frequency (Fig. 5a-c).

Secondly, we aimed to link clade-specific variation in resonant mechanics (i.e. proximity to resonance and *N*) to clade-specific behavioral and energetic requirements. We developed a new metric of aerodynamic efficiency that takes into account both *N* and 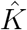 based upon the original work of Weis-Fogh. We found that silkmoths operate very close to undamped resonance (Fig. 5a-c) and have a lower *N* on average than hawkmoths (Fig. 7a), which results in a shallower resonance curve and higher aerodynamic efficiency (Fig. 7a,c). In contrast, hawkmoths operate farther from resonance than silkmoths and their larger *N* places them in a location in efficiency space with a steep gradient. Thus, *N* appears to be an important determinant of aerodynamic efficiency, dictating the degree to which off-resonance behavior incurs an energetic penalty. Indeed, in this region of efficiency space, a difference in *N* by 1 can reduce efficiency by nearly 20% (Fig. 7a). This result is particularly interesting in light of the fact that silkmoths do not feed as adults, as they lack functional mouthparts (*24*). As such, they are significantly more nutrient-limited over their life history, which may have led to selection for higher aerodynamic efficiency as a means of increasing their viable reproductive period (*21*). Hawkmoths instead may take advantage of their above-resonance wingbeats to more easily modulate wingbeat frequency to maneuver while tracking flowers (*7*), while maintaining sustainable energetics by virtue of a sharper resonance curve.

### 4.1 Constraints on wingbeat frequency-scaling of thorax and wing properties

Our transmission ratio scaling results (Fig. 3g-i) illustrate how muscle physiology may place a fundamental constraint on resonant mechanics. It is generally the case that faster movements are generated by smaller muscle strains. This allows muscles to operate on a narrow, more favorable location on their length-tension curve and produce higher forces at lower velocities, thus producing larger amounts of power (*20*). As such, our data suggests that these inherent muscle properties that mandate an increasing transmission ratio with frequency may be a more important constraint on thorax mechanics than any potential efficiency benefit from a stiffer thorax. Indeed, both hawkmoths and silkmoths already offset at least 50% of their inertial power costs with elastic energy storage over a range of wingbeat frequencies (Fig. 4c), suggesting that there might not be a large advantage to tuning thorax stiffness more precisely.

Biomechanical constraints imposed by thorax geometry and material may provide additional context for the invariance of thorax stiffness in moths. The thoracic shell is a highly intricate structure, with a shape that localizes strain energy in certain regions of exoskeleton (*12, 40*). Shape, rather than material, likely determines bulk stiffness, similar to how curvature affects stiffness of the human foot arch (*41*). Despite variation in thorax shape and size between moths of different species, we show that bulk stiffness remains the same (Fig. 3a-c), which is ultimately the stiffness that muscles encounter when actuating flight. As such, it may be very difficult to precisely tune thorax stiffness over evolutionary time without interfering with its functionality or structural integrity. In addition, it seems inefficient to modulate resonant frequency via thorax stiffness. Since resonant frequency is proportional only to the square root of stiffness, very large stiffness changes would be necessary to manifest in substantial resonant frequency variation across species. Such large stiffness variation may have negative consequences for the animal, like prohibitively high impedance for the flight muscles during takeoff. This does not preclude the hypothesized action of steering muscles to modulate stiffness within an animal over short timescales, as frequency modulation ranges for insects like *Manduca* are small compared to inter-species frequency differences (*7*). Finally, since aerodynamic and inertial power requirements scale sharply with frequency, a large amount of flight muscle in proportion to body size is required to drive flight in faster organisms (*42*). A need to pack as much power muscle as possible into the thorax may restrict any geometry-driven stiffness variation in the thorax over evolutionary time. Comparative analysis of muscle and thorax morphology across this group may shed light on how muscle constrains thorax properties.

Despite stiffness being nearly invariant and transmission ratio increasing with wingbeat frequency, we still find a proportional relationship between resonant frequency and wingbeat frequency. This is reflective of the importance of wing inertia in dictating an insect’s resonant mechanics. Wing inertia-frequency scaling is strong enough to compensate for the increase in transmission ratio that is required to generate high frequency wingbeats. Wing inertia in moths is a combination of wing mass and shape, the latter being a trait that varies extensively across Lepidopterans (*22*). We propose that maintaining favorable resonant mechanics is an additional pressure in the evolutionary tuning of moth wing shape and size, counteracting the effects of increasing transmission ratio with wingbeat frequency. In cold-hardy geometrid moths, low wing inertia has been shown to be the primary driver of reduced flight power costs that enable flight at extremely low body temperatures (*43*). Thus, wing inertia appears to be a particularly efficient ‘knob’ by which to tune flight energetics over evolutionary time in moths.

### 4.2 Multiple resonant peaks, non-linearity, and band-type resonance

We apply a simple model of a single resonance peak that is grounded in recent detailed work on *Manduca* (*5*). However, any conclusion that an animal is or is not at resonance depends on the precise definition of resonance being used. We provide comparisons of moth wingbeat frequency to both damped (displacement) and undamped (velocity) resonance, but note that the aerodynamic efficiency depends only on proximity to undamped resonance 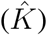. Undamped resonance is the frequency at which no dissipation is required by the muscle to drive flight, and represents the frequency at which inertial and elastic costs are instantaneously balanced at every point in a wingstroke (Fig. 6b,e). Most recent studies of insect resonance have been concerned with damped resonance, which maximizes wingbeat amplitude for a given input muscle force. Thus, operating at damped resonance may also be optimal, just by the different criterion of maximal wingbeat amplitude as opposed to aerodynamic efficiency. Our work suggests that investigation of multiple resonant frequencies and efficiency metrics may be necessary to fully contextualize an organisms’ preferred movement frequency.

Indeed, recent work by Pons et al. have demonstrated multiple distinct resonance frequencies that exist in the presence of thoracic nonlinearities in the flight motor (*17,44*). Such nonlin- earites in hawkmoths are likely small (*12*), but may become more significant at the high strains experienced by silkmoth thoraces. Similarly, we do not explicitly include the effects of active muscle stiffness. While muscle itself can store and return energy (*45*), active muscle stiffness is low compared to thorax stiffness in *Manduca* (*29*) and thus does not contribute highly to resonance. This is likely not the case in some small flying insects, like flies, where muscle stiffness is the dominant stiffness in the thorax (*18*). We do not know whether active muscle stiffness introduces a more significant non-linearity in high-strain silkmoths. In addition, we do not explicitly model series elasticity in the wing hinge, on the grounds that such effects are likely extremely small in *Manduca* (*5, 46*). Inclusion of substantial series elasticity in the wing hinge would primarily serve to widen the resonant peak, increasing the allowable frequencies of operation with minimal loss of efficiency (*17, 44*). In summary of all available evidence, wing hinge compliance and thoracic nonlinearity likely do not strongly influence resonance in bombycoids, but are increasingly important at the scale of *Drosophila* and smaller.

### 4.3 Insect flight resonance beyond moths

We focus on Bombycoidea as a model clade for studying resonant mechanics in closely related species against the backdrop of significant wing morphological and behavioural diversity.

However, we highlight a number of general principles that may apply broadly to other groups of insects, and areas for further comparative study. Increasing transmission ratio with wingbeat frequency is a general feature of flapping systems (*20*), and imposes a constraint on resonance in any clade that varies in wingbeat frequency. Any group of insects that vary in size will have to work against transmission ratio scaling to achieve resonance tuning. We show that in bombycoids, wing inertia scaling is strong enough to overcome the transmission ratio (Fig. 3i, Fig. 4a, Fig. 5a), but in other clades with more geometrically similar wings across body sizes, this may not be the case. Taking advantage of large intraspecific variation in insects like bees and studying resonant properties at an individual level may shed light on whether inertia is the main driver of resonance tuning more generally (*47*).

Many clades of insects such as Coleoptera, Hymenoptera, and Diptera, do not control their wings with a time-periodic nervous system signal, but instead actuate flight via antagonistic stretch-activated flight muscles (*48, 49*). In so-called asynchronous insects, wingbeat frequency is emergent so they are often thought to be flapping at resonance by definition, and have little neural control over their wingbeat frequency. Even so, bumblebees are capable of buzzing at multiple discrete frequencies that correspond to different behaviors such as thermogenesis, communication, buzz pollination, and flight (*50*). Unlike hawkmoths which can modulate frequency by neural activity (*7*), bees most likely achieve different wingbeat frequencies by modulating resonant properties of their thorax with steering muscles or transmission ratio via changing wing deployment. If insects like bees are operating close to resonance, our results suggest that they can accommodate a larger *N* while maintaining higher aerodynamic efficiency. Estimates from bees suggest *N >* 6, which would incur a large efficiency loss if they were sufficiently off of resonance (Fig. 7a). Alternatively, bees may endure this loss while maintaining moderate elastic energy storage in order to modulate frequency more widely, suggesting that frequency control may be at least as important as power and efficiency in insect flight.

Our work highlights the multivariate, often conflicting demands on the musculoskeletal systems of animals that utilize fast, oscillatory locomotion. Unlike commonly studied spring-driven ballistic animal movements (*51*), flapping insects must negotiate challenges associated with power production, dissipation, and control often on a wingstroke-to-wingstroke basis. Similarly, myriad terrestrial animals take advantage of resonant mechanics to improve locomotor efficiency by cyclically storing energy in tendons or apodemes (*3*). Resonance tuning, while an elegant explanation for insects’ preferred frequency of movement, requires a particular combination of thoracic spring, muscle physiology, wing transmission, and wing shape properties. Each of these components serves multiple functions and may vary counter to the expectation from resonance, such as in the case of wing transmission. In the case of bombycoid moths, wing inertia appears to be the primary knob by which resonance tuning is achieved. But even though resonance frequency scales with wingbeat frequency, Bombycoids still operate somewhat off of resonance, both with and without damping. Tradeoffs between proximity to resonance and *N* allow both hawkmoths and silkmoths to fly with feasible aerodynamic efficiency while operating within the constraints imposed by their muscle, thorax morphology, and behavior.

## Acknowledgments

This work was supported by US National Science Foundation RAISE grant no. IOS-2100858 to S.S. and N.G. and 1554790 (MPS-PoLS) and a Dunn Family Professorship to S.S. as well as the US National Science Foundation Physics of Living Systems SAVI student research network (GT node grant no. 1205878). Raw data are available from the Georgia Tech Digital Repository at: https://hdl.handle.net/1853/72950

## Author Contributions

E.S.W., J.E.L., N.G., S.S., B.A., and U.S. conceived of the study. E.S.W. and M.H. ran experiments. E.S.W. analyzed data, and prepared the figures. All authors wrote and edited the manuscript.

## Notes

### Competing Interest Statement

The authors have declared no competing interest.

