## Supplementary Information for "Moth resonant mechanics are tuned to wingbeat frequency and energetic demands"

#### 1 Calculation of Weis-Fogh number from flight power

The Weis-Fogh number ($N$) was originally defined as the ratio of peak aerodynamic and inertial torques over a wingstroke (*1*):

$$N = \frac{\max\left(\tau_{inertial}\right)}{\max\left(\tau_{aero}\right)} = \frac{I}{\Gamma\phi_o} \tag{1}$$

While estimates of $N$ using this equation are reasonably accurate when one knows all required information for an insect of interest, it can be challenging to apply Eq. 1 comparatively. Either one must know peak torques from a model (preferably one that incorporates 3D kinematics and/or computational fluid dynamics), or one must know species-specific average drag coefficients and locations of the center of pressure on the wing. Peak torque estimates are not widely reported and wind-tunnel drag coefficient data for many species is not available.

Instead, we use more commonly-reported mean cycle-averaged aerodynamic ($\bar{P}_{aero}$) and inertial powers ($\bar{P}_{inertial}$) to compute $N$. We derive an expression for $N$ that depends on the ratio of these powers using the equation of motion in the main text (main text Eq. 1). Then, we use power estimates for each species from a recent blade-element model that incorporates species-specific wingbeat kinematics to estimate $N$ comparatively.

First, we must compute average inertial and aerodynamic power from main text Eq. 1 by integrating the instantaneous power, defined as torque multiplied by angular velocity. Our integration bounds will be the first quarter-stroke ($t = 0$ to $t = \pi/2\omega$), since this is the region during which both aerodynamic and inertial powers are positive. Since we seek the ratio of the mean absolute value of these powers, the result for a quarter stroke will be identical to that for a full stroke and much easier to compute.

The inertial power can be written as:

$$\bar{P}_{inertial} = \frac{\pi}{2\omega}\int_0^{\pi/2\omega} I\ddot{\phi}(t)\dot{\phi}(t)dt = \frac{I\phi_o^2\omega^3\pi}{2\omega}\int_0^{\pi/2\omega} \sin(2\omega t)dt = \frac{1}{2}I\phi_o^2\omega^2 \tag{2}$$

The aerodynamic power can be written as:

$$\bar{P}_{aero} = \frac{\pi}{2\omega}\int_0^{\pi/2\omega} \Gamma|\dot{\phi}(t)|(\dot{\phi}(t))^2dt = \frac{\Gamma\phi_o^3\omega^3\pi}{2\omega}\int_0^{\pi/2\omega} \cos^2(\omega t)|\cos(\omega t)|dt = \frac{2}{3}\Gamma\phi_o^3\omega^2 \tag{3}$$

Taking the ratio of Eqs. 2 and 3 and substituting in Eq. 1, we arrive at the final expression for $N$ in terms of $\bar{P}_{inertial}$ and $\bar{P}_{aero}$.

$$N = \frac{4}{3}\frac{\bar{P}_{inertial}}{\bar{P}_{aero}} \tag{4}$$

$\bar{P}_{inertial}$ and $\bar{P}_{aero}$ are then computed for each species using the same model of Aiello et al. 2021 (*2*). Aerodynamic power includes contributions from induced, profile, and parasitic power. See the citation and citation supplement for details.

### 2 Calculation of aerodynamic efficiency

As discussed in the main text, we derive a new aerodynamic efficiency metric that applies to flapping insects that are not necessarily flapping at resonance. We do this by following the original logic of Weis-Fogh, defining efficiency as the ratio of positive aerodynamic work over a cycle to total positive work (sum of inertial, aerodynamic, and elastic contributions) (*1*). Following the work of Lynch et al., we begin with the non-dimensional torques as a function of non-dimensional wing angle (*3*):

$$\hat{\tau}_{aero}(t) = 1 - (\hat{\phi}(t))^2 \tag{5}$$

$$\hat{\tau}_{inertial}(t) = -N\hat{\phi}(t) \tag{6}$$

$$\hat{\tau}_{elastic}(t) = \hat{K}N\hat{\phi}(t) \tag{7}$$

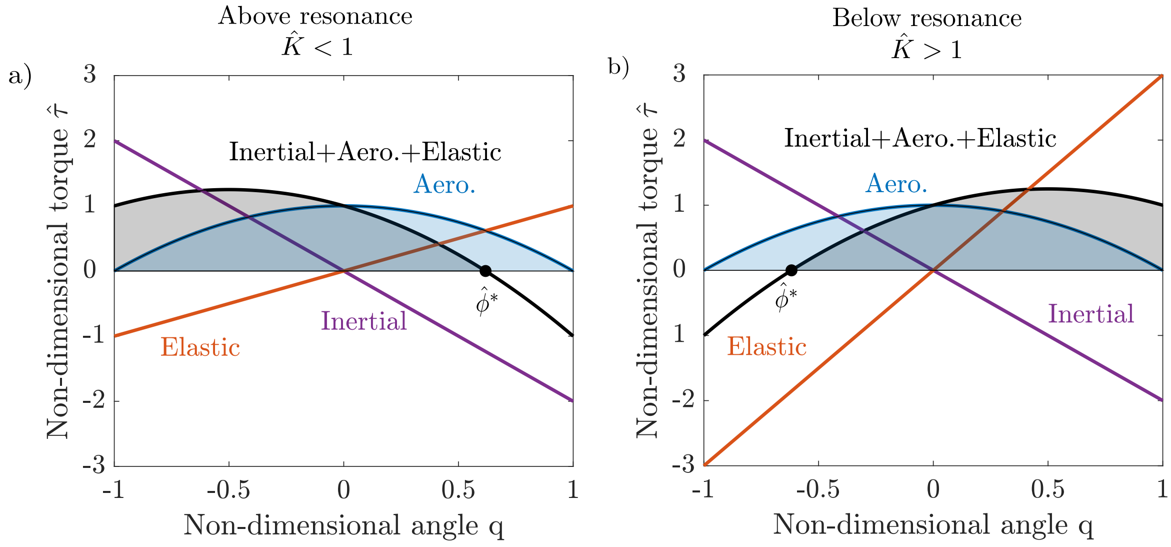


Figure 1: a). Non-dimensional torques as a function of wing angle in the above-resonance case, with the integration bound $\hat{\phi}^*$ notated. b). Non-dimensional torques as a function of wing angle in the below-resonance case, with the integration bound $\hat{\phi}^*$ notated. Shaded blue and black areas correspond to aerodynamic work and total positive work respectively.

Aerodynamic efficiency is then defined by the following integral expression, where bounds of integration are chosen to ensure only positive work is being considered.

$$\eta = 100 \frac{\int_+ \hat{\tau}_{aero}(t) d\hat{\phi}}{\int_+ \hat{\tau}_{inertial}(t) + \hat{\tau}_{elastic}(t) + \hat{\tau}_{aero}(t) d\hat{\phi}} \tag{8}$$

From here on, $\hat{\tau}$ and $\hat{\phi}$ are implied to be functions of $t$ for ease of notation. Since $\hat{\tau}_{aero}$ is nonnegative over the domain $\hat{\phi} \in [-1, 1]$, we can integrate equation 14 directly over this domain:

$$\int_{-1}^{1} (1 - \hat{\phi}^2) d\hat{\phi} = 4/3 \tag{9}$$

Note that if $\hat{\tau}_{inertial} = \hat{\tau}_{elastic}$, efficiency is always $100\%$, which is the case when the insect is flapping at undamped resonance, $\hat{K} = 1$. So, the denominator integral can be decomposed into two cases: when $\hat{K} < 1$, and when $\hat{K} > 1$. In each of these cases, the sum of inertial, elastic, and aerodynamic torques is not positive across the whole domain of integration. We can compute the $\hat{\phi}$ where the sum of torques crosses zero by finding the zeros of the following two quadratic equations for the $\hat{K} < 1$ and $\hat{K} > 1$ conditions respectively:

$$1 - \hat{\phi}^2 + N(1 - \hat{K})\hat{\phi} = 0 \tag{10}$$

$$1 - \hat{\phi}^2 + N(\hat{K} - 1)\hat{\phi} = 0 \tag{11}$$

If we call the zeros of these equations $\hat{\phi}^*_{\hat{K}<1}$ and $\hat{\phi}^*_{\hat{K}>1}$, we can express the denominator of Eq. 17 piecewise by the following two integrals:

$$\int_{+} \hat{\tau}_{inertial} + \hat{\tau}_{elastic} + \hat{\tau}_{aero} d\hat{\phi} = \left\{ \begin{array}{ll} \int_{-1}^{\hat{\phi}^*_{\hat{K}<1}} 1 - \hat{\phi}^2 + N(1 - \hat{K})\hat{\phi} d\hat{\phi}, & \text{if } \hat{K} < 1 \\ \int_{\hat{\phi}^*_{\hat{K}>1}}^{1} 1 - \hat{\phi}^2 + N(\hat{K} - 1)\hat{\phi} d\hat{\phi}, & \text{if } \hat{K} > 1 \end{array} \right\} \tag{12}$$

Combining the result of Eq. 18 with Eq. 21, one can easily compute $\eta$ from Eq. 17 with a computer algebra program like Mathematica.
